# Efficient Construction of a Complete Index for Pan-Genomics Read Alignment

**DOI:** 10.1101/472423

**Authors:** Alan Kuhnle, Taher Mun, Christina Boucher, Travis Gagie, Ben Langmead, Giovanni Manzini

## Abstract

While short read aligners, which predominantly use the FM-index, are able to easily index one or a few human genomes, they do not scale well to indexing databases containing thousands of genomes. To understand why, it helps to examine the main components of the FM-index in more detail, which is a rank data structure over the Burrows-Wheeler Transform (BWT) of the string that will allow us to find the interval in the string’s suffix array (SA) containing pointers to starting positions of occurrences of a given pattern; second, a sample of the SA that — when used with the rank data structure — allows us access the SA. The rank data structure can be kept small even for large genomic databases, by run-length compressing the BWT, but until recently there was no means known to keep the SA sample small without greatly slowing down access to the SA. Now that Gagie et al. (SODA 2018) have defined an SA sample that takes about the same space as the run-length compressed BWT — we have the design for efficient FM-indexes of genomic databases but are faced with the problem of building them. In 2018 we showed how to build the BWT of large genomic databases efficiently (WABI 2018) but the problem of building Gagie et al.’s SA sample efficiently was left open. We compare our approach to state-of-the-art methods for constructing the SA sample, and demonstrate that it is the fastest and most space-efficient method on highly repetitive genomic databases. Lastly, we apply our method for indexing partial and whole human genomes, and show that it improves over Bowtie with respect to both memory and time.

**Availability:** We note that the implementation of our methods can be found here: https://github.com/alshai/r-index.

## 1 Introduction

The FM-index, which is a compressed subsequence index based on Burrows Wheeler transform (BWT), is the primary data structure majority of short read aligners — including Bowtie [19], BWA [13] and SOAP2 [21]. These aligners build a FM-index based data structure of sequences from a given genomic database, and then use the index to perform queries that find approximate matches of sequences to the database. And while these methods can easily index one or a few human genomes, they do not scale well to indexing the databases of thousands of genomes. This is problematic in analysis of the data produced by consortium projects, which routinely have several thousand genomes.

In this paper, we address this need by introducing and implementing an algorithm for efficiently constructing the FM-index. This will allow for the FM-index construction to scale to larger sets of genomes. To understand the challenge and solution behind our method, it helps to examine the two principal components of the FM-index: first a rank data structure over the BWT of the string that enables us to find the interval in the string’s suffix array (SA) containing pointers to starting positions of occurrences of a given pattern (and, thus, compute how many such occurrences there are); second, a sample of the SA that, when used with the rank data structure, allows us access the SA (so we can list those starting positions). Searching with an FM-index can be summarized as follows: starting with the empty suffix, for each proper suffix of the given pattern we use rank queries at the ends of the BWT interval containing the characters immediately preceding occurrences of that suffix in the string, to compute the interval containing the characters immediately preceding occurrences of the suffix of length 1 greater; when we have the interval containing the characters immediately preceding occurrences of the whole pattern, we use a SA sample to list the contexts of the corresponding interval in the SA, which are the locations of those occurrences.

Although it is possible to use a compressed implementation of the rank data structure that does not become much slower or larger even for thousands of genomes, the same cannot be said for the SA sample. The product of the size and the access time must be at least linear in the length of the string for the standard SA sample. This implies that the FM-index will become much slower and/or much larger as the number of genomes in the databases grows significantly. This bottleneck has forced researchers to consider variations of FM-indexes adapted for massive genomic datasets, such as Valenzuela et al.’s pan-genomic index [33] or Garrison et al.’s variation graphs [7]. Some of these proposals use elements of the FM-index, but all deviate in substantial ways from the description above. Not only does this mean they lack the FM-index’s long and successful track record, it also means they usually do not give us the BWT intervals for all the suffixes as we search (whose lengths are the suffixes’ frequencies, and thus a tightening sequence of upper bounds on the whole pattern’s frequency), nor even the final interval in the suffix array (which is an important input in other string processing tasks).

Recently, Gagie, Navarro and Prezza [11] proposed a different approach to SA sampling, that takes space proportional to that of the compressed rank data structure while still allowing reasonable access times. While their result yields a potentially practical FM-index on massive databases, it does not directly lead to a solution since the problem of how to efficiently construct the BWT and SA sample remained open. In a direction toward to fully realizing the theoretical result of Gagie et al. [11], Boucher et al. [2] showed how to build the BWT of large genomic databases efficiently. We refer to this construction as *prefix-free parsing*. It takes as input string *S*, and in one-pass generates a dictionary and a parse of S with the property that the BWT can be constructed from dictionary and parse using workspace proportional to their total size and *O*(|*S*|) time. Yet, the resulting index of Boucher et al. [2] has no SA sample, and therefore, only supports counting and not locating. This makes this index not directly applicable to many bioinformatic applications, such as sequence alignment.

### Our contributions

In this paper, we present a solution for building the FM-index^1^ for very large datasets by showing that we can build the BWT and Gagie et al.’s SA sample together in roughly the same time and memory needed to construct the BWT alone. We note that this algorithm is also based on prefix-free parsing. Thus, we begin by describing how to construct the BWT from the prefix-free parse, and then show that it can be modified to build the SA sample in addition to the BWT in roughly the same time and space. We implement this approach, and refer to the resulting implementation as `bigbwt`. We compare it to state-of-the-art methods for constructing the SA sample, and demonstrate that `bigbwt` currently the fastest and most space-efficient method for constructing the SA sample on large genomic databases.

Next, we demonstrate the applicability of our method to short read alignment. In particular, we compare the memory and time needed by our method to build an index for collections of chromosome 19 with that of Bowtie. Through these experiments, we show that Bowtie was unable to build indexes for our largest collections (500 or more) because it exhausted memory, whereas our method was able to build indexes up to 1,000 chromosome 19s (and likely beyond). At 250 chromosome 19 sequences, the our method required only about 2% of the time and 6% the peak memory of Bowtie’s. Lastly, we demonstrate that it is possible to index collections of whole human genome assemblies with sub-linear scaling as the size of the collection grows.

### Related work

The development of methods for building and the FM-index on large datasets is closely related to the development short-read aligners for pan-genomics — an area where there is growing interest [27,5,12]. Here, we briefly describe some previous approaches to this problem and detail its connection to the work in this paper. We note that majority of pan-genomic aligners requiring building the FM-index for a population of genomes and thus, can increase proficiency by using the methods described in this paper.

GenomeMapper [27], the method of Danek et al. [5], and GCSA [29] represent the genomes in a population as a graph, and then reduce the alignment problem to finding a path within the graph. Hence, these methods require all possible paths to be identified, which is exponential in the worst case. Some of these methods — such as the GCSA — use the FM-index to store and query the graph and could capitalize on our approach by building the index in the manner described here. Another set of approaches [24,8,12,32] consider the reference pan-genome as the concatenation of individual genomes and exploits redundancy by using a compressed index. The hybrid index [8] operates on a Lempel-Ziv compression of the reference pan-genome. An input parameter *M* sets the maximum length of reads that can be aligned; the parameter *M* has a large impact on the final size of the index. For this reason, the hybrid index is suitable for short-read alignment only, although there have been recent heuristic modifications to allow longer alignments [9]. In contrast, the *r*-index, of which we provide an implementation in this work, has no such length limitation. The most recent implementation of the hybrid index is CHIC [33]. Although CHIC has support for counting multiple occurrences of a pattern within a genomic database, it is an expensive operation, namely *O*(*ℓ* log log *n*), where *ℓ* is the number of occurrences in the databases and *n* is the length of the database. However, the *r*-index is capable of counting all occurrences of a pattern of length m in *O*(*m*) time up to polylog factors. There are a number of other approaches building off the hybrid index or similar ideas [5,34]; for an extended discussion, we refer the reader to the survey of Gagie and Puglisi [12].

Finally, a third set of approaches [14,23] attempts to encode variants within a single reference genome. BWBBLE by Huang et al. [14] follows this by supplementing the alphabet to indicate if multiple variants occur at a single location. This approach does not support counting of the number of variants matching a specific alignment; also, it suffers from memory blow-up when larger structural variations occur.

## 2 Background

### 2.1 BWT and FM indexes

Consider a string *S* of length *n* from a totally ordered alphabet *Σ*, such that the last character of *S* is lexicographically less than any other character in *S*. Let *F* be the list of *S*’s characters sorted lexicographically by the suffixes starting at those characters, and let *L* be the list of *S*’s characters sorted lexicographically by the suffixes starting immediately after those characters. The list L is termed the Burrows-Wheeler Transform [3] of *S* and denoted BWT. If *S*[*i*] is in position *p* in *F* then *S*[*i* − 1] is in position *p* in *L*. Moreover, if *S*[*i*] = *S*[*j*] then *S*[*i*] and *S*[*j*] have the same relative order in both lists; otherwise, their relative order in *F* is the same as their lexicographic order. This means that if *S*[*i*] is in position *p* in *L* then, assuming arrays are indexed from 0 and ≺ denotes lexicographic precedence, in *F* it is in position *j_i_* = |{*h*: *S*[*h*] ≺ *S*[*i*]}| + |{*h*: *L*[*h*] = *S*[*i*], *h* ≤ *p*}| − 1. The mapping *i* ↦ *j_i_* is termed the LF mapping. Finally, notice that the last character in *S* always appears first in *L*. By repeated application of the LF mapping, we can invert the BWT, that is, recover *S* from *L*. Formally, the *suffix array* SA of the string *S* is an array such that entry *i* is the starting position in *S* of the *i*th largest suffix in lexicographical order. The above definition of the BWT is equivalent to the following:

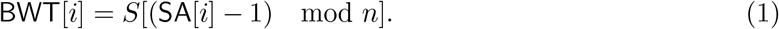

The BWT was introduced as an aid to data compression: it moves characters followed by similar contexts together and thus makes many strings encountered in practice locally homogeneous and easily compressible. Ferragina and Manzini [10] showed how the BWT may be used for *indexing* a string *S*: given a pattern *P* of length *m* < *n*, find the number and location of all occurrences of *P* within *S*. If we know the range BWT(*S*)[*i*..*j*] occupied by characters immediately preceding occurrences of a pattern *Q* in *S*, then we can compute the range BWT(*S*)[*i*′..*j*′] occupied by characters immediately preceding occurrences of *cQ* in *S*, for any character *c* ∈ *Σ*, since

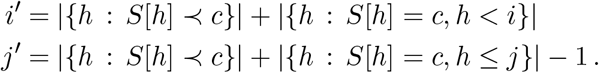

Notice *j*′ − *i*′ + 1 is the number of occurrences of *cQ* in *S*. The essential components of an FM-index for *S* are, first, an array storing |{*h*: *S*[*h*] ≺ *c*}| for each character c and, second, a rank data structure for BWT that quickly tells us how often any given character occurs up to any given position^2^. To be able to locate the occurrences of patterns in *S* (in addition to just counting them), the FM-index uses a sampled^3^ suffix array of *S* and a bit vector indicating the positions in BWT of the characters preceding the sampled suffixes.

### 2.2 Prefix-free parsing

Next, we give an overview of prefix-free parsing, which produces a dictionary 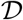 and a parse 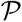 by sliding a window of fixed width through the input string *S*. We refer the reader to Boucher et al. [2] for the formal proofs and Section 3.1 for the algorithmic details. A rolling hash function identifies when substrings are parsed into elements of a dictionary, which is a set of substrings of *S*. Intuitively, for a repetitive string, the same dictionary phrases will be encountered frequently.

We now formally define the dictionary 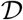 and parse 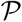. Given a string^4^ *S* of length *n*, window size *w* ∈ ℕ and modulus *p* ∈ ℕ, we construct the dictionary 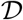 of substrings of *S* and the parse 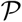 as follows. We let *f* be a hash function on strings of length *w*, and let 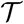 be the sequence of substrings *W* = *S*[*s, s* + *w* − 1] such that *f*(*W*) = 0 mod *p* or *W* = *S*[0, *w* − 1] or *W* = *S*[*n* − *w* + 1, *n*], ordered by initial position in *S*; let 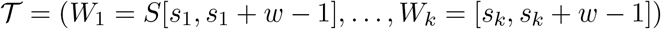. By construction the strings

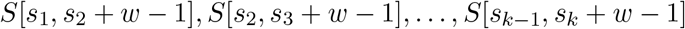

form a parsing of *S* in which each pair of consecutive strings *S*[*s_i_, s*_*i*+1_ + *w* − 1] and *S*[*s*_*i*+1_, *s*_*i*+2_ + *w* − 1] overlaps by exactly *w* characters. We define 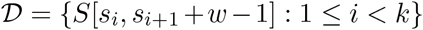; that is, 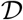 consists of the set of the unique substrings *s* of *S* such that |*s*| > *w* and the first and last *w* characters of *s* form consecutive elements in 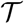. If *S* has many repetitions we expect that 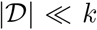. With a little abuse of notation we define the parsing 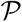 as the sequence of lexicographic ranks of substrings in 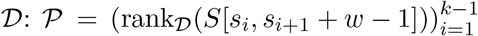. The parse 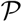 indicates how *S* may be reconstructed using elements of 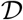. The dictionary 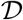 and parse 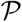 may be constructed in one pass over *S* in 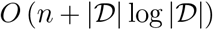 time if the hash function *f* can be computed in constant time.

### 2.3 r-index locating

Policriti and Prezza [26] showed that if we have stored SA[*k*] for each value *k* such that BWT[*k*] is the beginning or end of a run (i.e., a maximal non-empty unary substring) in BWT, and we know both the range BWT[*i..j*] occupied by characters immediately preceding occurrences of a pattern *Q* in *S* and the starting position of one of those occurrences of *Q*, then when we compute the range BWT[*i*′..*j*′] occupied by characters immediately preceding occurrences of *cQ* in *S*, we can also compute the starting position of one of those occurrences of *cQ*. Bannai et al [1] then showed that even if we have stored only SA[*k*] for each value *k* such that BWT[*k*] is the beginning of a run, then as long as we know SA[*i*] we can compute SA[*i*′].

Gagie, Navarro and Prezza [11] showed that if we have stored in a predecessor data structure SA[*k*] for each value *k* such that BWT[*k*] is the beginning of a run in BWT, with *φ*^−1^(SA[*k*]) = SA[*k* + 1] stored as satellite data, then given SA[*h*] we can compute SA[*h* + 1] in *O*(log log *n*) time as SA[*h* + 1] = *φ*^−1^(pred(SA[*h*])) + SA[*h*] − pred(SA[*h*]), where pred(·) is a query to the predecessor data structure. Combined with Bannai et al.’s result, this means that while finding the range BWT[*i..j*] occupied by characters immediately preceding occurrences of a pattern *Q*, we can also find SA[*i*] and then report SA[*i* + 1..*j*] in *O*((*j* − *i*) log log *n*)-time, that is, *O*(log log *n*)-time per occurrence.

Gagie et al. gave the name *r*-index to the index resulting from combining a rank data structure over the run-length compressed BWT with their SA sample, and Bannai et al. used the same name for their index. Since our index is an implementation of theirs, we keep this name; on the other hand, we do not apply it to indexes based on run-length compressed BWTs that have standard SA samples or no SA samples at all.

## 3 Methods

Here, we describe our algorithm for building the SA or the sampled SA from the prefix free parse of a input string *S*, which is used to build the *r*-index. We first review the algorithm from [2] for building the BWT of S from the prefix free parse. Next, we show how to modify this construction to compute the SA or the sampled SA along with the BWT.

### 3.1 Construction of BWT from prefix-free parse

We assume we are given a prefix-free parse of *S*[1..*n*] with window size *w* consisting of a dictionary 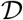 and a parse 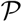. We represent the dictionary as a string 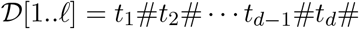 where *t_i_*’s are the dictionary phrases in lexicographic order and # is a unique separator. We assume we have computed the SA of 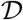, denoted by 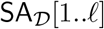 in the following, and the suffix array of 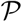, denoted 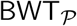, and the array Occ[1, *d*] such that Occ[*i*] stores the number of occurrences in the parse of the dictionary phrase *t_i_*. These preliminary computations take 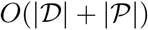 time.

By the properties of the prefix-free parsing, each suffix of *S* is prefixed by *exactly one* suffix *α* of a dictionary phrase *t_j_* with |*α*| > *w*. We call α the *representative prefix* of the suffix *S*[*i..n*]. From the uniqueness of the representative prefix we can partition *S*’s suffix array SA[1..*n*] into *k* ranges

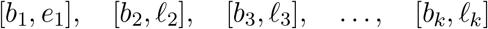

with *b*_1_ = 1, *b_i_* = *e_i_* + 1 for *i* = 2,…, *k*, and *e_k_* = *n*, such that for *i* = 1,…, *k* all suffixes

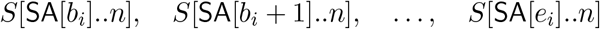

have the same representative prefix *α_i_*. By construction *α*_1_ ≺ *α*_2_ ≺ ⋯ ≺ *α_k_*.

By construction, any suffix 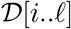 of the dictionary 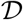 is also prefixed by the suffix of a dictionary phrase. For *j* = 1,…, *ℓ*, let *β_j_* denote the longest prefix of 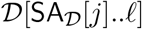 which is the suffix of a phrase (i.e. 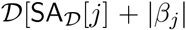). By construction the strings *β_j_*’s are lexicographically sorted *β*_1_ ≺ *β*_2_ ≺ ⋯ ≺ *β_ℓ_*. Clearly, if we compute *β*_1_, …, *β_ℓ_* and discard those such that |*β_j_*| ≤ *w*, the remaining *β_j_*’s will coincide with the representative prefixes *α_i_*’s. Since both *β_j_*’s and *α_i_*’s are lexicographically sorted, this procedure will generate the representative prefixes in the order *α*_1_, *α*_2_,…, *α_k_*. We note that more than one *β_j_* can be equal to some *α_i_* since different dictionary phrases can have the same suffix.

We scan 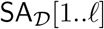, compute *β*_1_, … *β_ℓ_* and use these strings to find the representative prefixes. As soon as we generate an *α_i_* we compute and output the portion BWT[*b_i_, e_i_*] corresponding to the range [*b_i_, e_i_*] associated to *α_i_*. To implement the above strategy, assume there are exactly *k* entries in 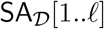 prefixed by *α_i_*. This means that there are *k* distinct dictionary phrases *t*_*i*_1__, *t*_*i*_2__,…, *t_i_k__* that end with *α_i_*. Hence, the range [*b_i_, e_i_*] contains 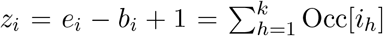 elements. To compute BWT[*b_i_, e_i_*] we need to: 1) find the symbol immediately preceding each occurrence of *α_i_* in *S*, and 2) find the lexicographic ordering of *S*’s suffixes prefixed by *α_i_*. We consider the latter problem first.

#### Computing the lexicographic ordering of suffixes

For *j* = 1,…, *z_i_* consider the *j*-th occurrence of *α_i_* in *S* and let *i_j_* denote the position in the parsing of *S* of the phrase ending with the *j*-th occurrence of *α_i_*. In other words, 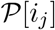 is a dictionary phrase ending with *α_i_* and *i*_1_ < *i*_2_ < ⋯ < *i_z_i__*. By the properties of 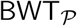 the lexicographic ordering of *S*’s suffixes prefixed by *α_i_* coincides with the ordering of the symbols 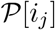 in 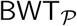. In other words, 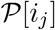 precedes 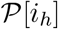 in 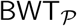 if and only if *S*’s suffix prefixed by the *j*-th occurrence of *α_i_* is lexicographically smaller than *S*’s suffix prefixed by the *h*-th occurrence of *α_i_*.

We could determine the desired lexicographic ordering by scanning 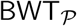 and noticing which entries coincide with one of the dictionary phrases *t*_*i*_1__,…, *t_i_k__* that end with *α_i_* but this would clearly be inefficient. Instead, for each dictionary phrase *t_i_* we maintain an array IL_*i*_ of length Occ[*i*] containing the indexes *j* such that 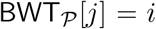. These sorts of “inverted lists” are computed at the beginning of the algorithm and replace the 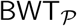 which can be discarded.

#### Finding the symbol preceding α_i_

Given a representative prefix *α_i_* from 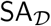 we retrieve the indexes *i*_1_,…, *i_k_* of the dictionary phrases *t*_*i*_1__,…, *t_i_k__* that end with *α_i_*. Then, we retrieve the inverted lists IL_*i*_1__,… IL*_i_k__* and we merge them obtaining the list of the *z_i_* positions *y*_1_ < *y*_2_ < ⋯ < *y_z_i__* such that BWT_*P*_[*y_j_*] is a dictionary phrase ending with *α_i_*. Such list implicitly provides the lexicographic order of *S*’s suffixes starting with *α_i_*.

To compute the BWT we need to retrieve the symbols preceding such occurrences of *α_i_*. If *α_i_ is not* a dictionary phrase, then *α_i_* is a proper suffix of the phrases *t*_*i*_1__,…, *t_i_k__* and the symbols preceding *α_i_* in *S* are those preceding *α_i_* in *t*_*i*_1__,…, *t_i_k__* that we can retrieve from 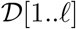 and 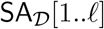. If *α_i_ coincides* with a dictionary phrase *t_j_*, then it cannot be a suffix of another phrase. Hence, the symbols preceding *α_i_* in *S* are those preceding *t_j_* in *S* that we store at the beginning of the algorithm in an auxiliary array PR_*j*_ along with the inverted list IL_*j*_.

### 3.2 Construction of SA and SA sample along with the BWT

We now show how to modify the above algorithm so that, along with BWT, it computes the full SA of *S* or the sampled SA consisting of the values SA[*s*_1_],…, SA[*s_r_*] and SA[*e*_1_],…, SA[*e_r_*], where *r* is the number of maximal non-empty runs in BWT and *s_i_* and *e_i_* are the starting and ending positions in BWT of the *i*-th such run, respectively. Note that if we compute the sampled SA the actual output will consist of *r* start-run pairs 〈*s_i_*, SA[*s_i_*]〉 and *r* end-run pairs 〈*e_i_*, SA[*e_i_*]〉 since the SA values alone are not enough for the construction of the *r*-index.

We solve both problems using the following strategy. Simultaneously to each entry BWT[*j*], we compute the corresponding entry SA[*j*]. Then, if we need the sampled SA, we compare BWT[*j* − 1] and BWT[*j*] and if they differ, we output the pair 〈*j* − 1, SA[*j* − 1]〉 among the end-runs and the pair 〈*j*, SA[*j*]〉 among the start-runs. To compute the SA entries, we only need *d* additional arrays EP_1_,… EP_*d*_ (one for each dictionary phrase), where |EP_*i*_| = |IL_*i*_| = Occ[*i*], and EP_*i*_[*j*] contains the ending position in *S* of the dictionary phrase which is in position IL_*i*_[*j*] of 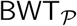.

Recall that in the above algorithm for each occurrence of a representative prefix *α_i_*, we compute the indexes *i*_1_,…, *i_k_* of the dictionary phrases *t*_*i*_1__,…, *t_i_k__* that end with *α_i_*. Then, we use the lists IL_*i*_1__,…, IL_*i_k_*_ to retrieve the positions of all the occurrences of *t*_*i*_1__,…, *t_i_k__* in 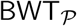, thus establishing the relative lexicographic order of the occurrences of the dictionary phrases ending with *α_i_*. To compute the corresponding SA entries, we need the starting position in *S* of each occurrence of *α_i_*. Since the ending position in *S* of the phrase with relative lexicographic rank IL_*i_h_*_[*j*] is EP_*i_h_*_[*j*], the corresponding SA entry is EP_*i_h_*_[*j*] − |*α_i_*| + 1. Hence, along with each BWT entry we obtain the corresponding SA entry which is saved to the output file if the full SA is needed, or further processed as described above if we need the sampled SA.

## 4 Time and memory usage for SA and SA sample construction

We compare the running time and memory usage of `bigbwt`^5^ with the following methods, which represent the current state-of-the-art.

`bwt2sa` Once the BWT has been computed, the SA or SA sample may be computed by applying the LF mapping to invert the BWT and the application of Eq. 1. Therefore, as a baseline, we use `bigbwt` to construct the BWT only, as in Boucher *et al*. [2]; next, we load the BWT as a Huffman-compressed string with access, rank, and select support to compute the LF mapping. We step backwards through the BWT and compute the entries of the SA in non-consecutive order. Finally, these entries are sorted in external memory to produce the SA or SA sample. This method may be parallelized when the input consists of multiple strings by stepping backwards from the end of each string in parallel.

`pSAscan` A second baseline is to compute the SA directly from the input; for this computation, we use the external-memory algorithm `pSAscan` [17], with available memory set to the memory required by `bigbwt` on the specific input; with the ratio of memory to input size obtained from `bigbwt, pSAscan` is the current state-of-the-art method to compute the SA. Once `pSAscan` has computed the full SA, the SA sample may be constructed by loading the input text T into memory, streaming the SA from the disk, and the application of Eq. 1 to detect run boundaries. We denote this method of computing the SA sample by `pSAscan+`.

We compared the performance of all the methods on two datasets: (1) Salmonella genomes obtained from GenomeTrakr [31]; and (2) chromosome 19 haplotypes derived from the 1000 Genomes Project phase 3 data [4]. The Salmonella strains were downloaded from NCBI (NCBI BioProject PRJNA183844) and preprocessed by assembling each individual sample with IDBA-UD [25] and counting *k*-mers (*k*=32) using KMC [6]. We modified IDBA by setting kMaxShortSequence to 1024 per public advice from the author to accommodate the longer paired end reads that modern sequencers produce. We sorted the full set of samples by the size of their *k*-mer counts and selected 1,000 samples about the median. This avoids exceptionally short assemblies, which may be due to low read coverage, and exceptionally long assemblies which may be due to contamination.

Next, we downloaded and preprocessed a collection of chromosome 19 haplotypes from 1000 Genomes Project. Chromosome 19 is 58 million base pairs in length and makes up around 1.9% of the total human genome sequence. Each sequence was derived by using the `bcftools consensus` tool to combine the haplotype-specific (maternal or paternal) variant calls for an individual in the 1KG project with the chr19 sequence in the GRCH37 human reference, producing a FASTA record per sequence. All DNA characters besides A, C, G, T and N were removed from the sequences before construction.

We performed all experiments in this section on a machine with Intel(R) Xeon(R) CPU E5-2680 v2 @ 2.80GHz and 324 GB RAM. We measured running time and peak memory footprint using `/usr/bin/time -v`, with peak memory footprint captured by the `Maximum resident set size (kbytes)` field and running time by the `User Time` and `System Time` field.

We witnessed that the running time of each method to construct the full SA is shown in Figs. 1(a) – 1(c). On both the Salmonella and chr19 datasets, `bigbwt` ran the fastest, often by more than an order of magnitude. In Fig. 1(d), we show the peak memory usage of `bigbwt` as a function of input size. Empirically, the peak memory usage was sublinear in input size, especially on the chr19 data, which exhibited a high degree of repetition. Despite the higher diversity of the Salmonella genomes, `bigbwt` remained space-efficient and the fastest method for construction of the full SA. Furthermore, we found qualitatively similar results for construction of the SA sample, shown in Fig. 2. Similar to the results on full SA construction, `bigbwt` outperformed both baseline methods and exhibited sublinear memory scaling on both types of databases.

**Fig. 1:**
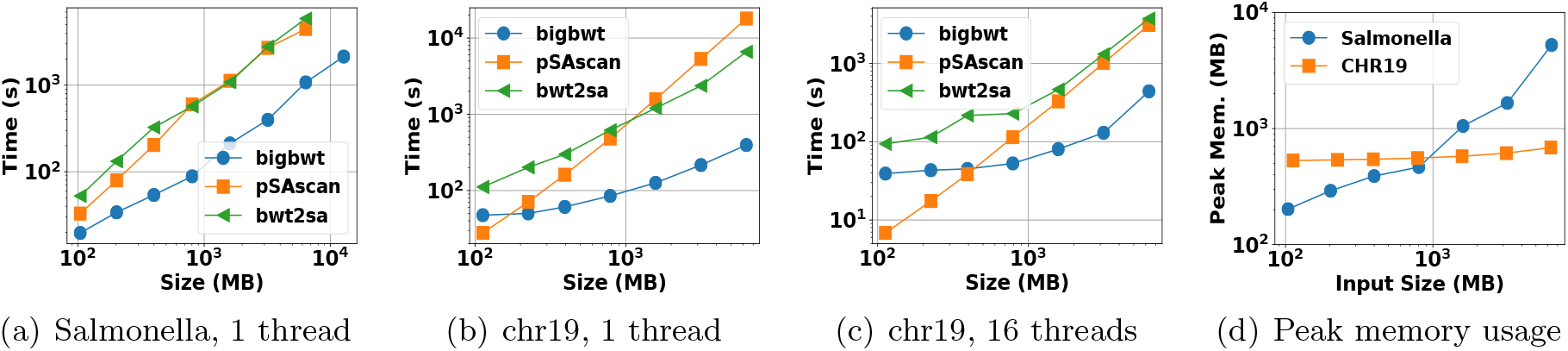
Runtime and peak memory usage for construction of full SA.

**Fig. 2:**
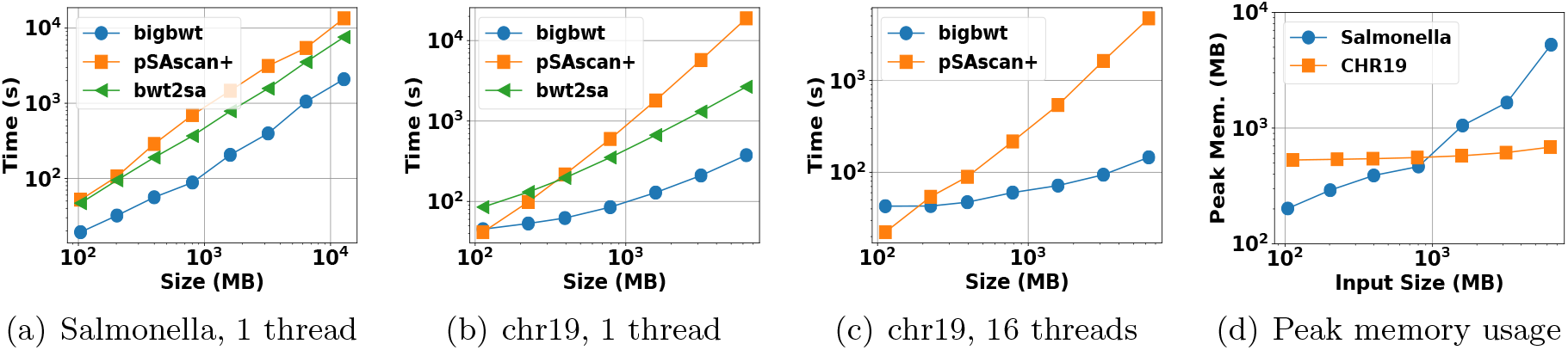
Runtime and peak memory usage for construction of SA sample.

## 5 Application to many human genome sequences

We studied how the *r*-index scales to repetitive texts consisting of many similar genomic sequences. Since an ultimate goal is to improve read alignment, we benchmark against Bowtie (version 1.2.2) [19]. We ran Bowtie with the `-v 0` and `--norc` options; `-v 0` disables approximate matching, while `--norc` causes Bowtie (like *r*-index) to perform the locate query with respect to the query sequence only and not its reverse complement.

### 5.1 Indexing chromosome 19s

We performed our experiments on collections of one or more versions of chromosome 19. These versions were obtained from 1000 Genomes Project haplotypes in the manner described in the previous section. We used 10 collections of chromosome 19 haplotypes, containing 1, 2, 10, 30, 50, 100, 250, 500, and 1000 sequences, respectively. Each collection is a superset of the previous. Again, all DNA characters besides A, C, G, T and N were removed from the sequences before construction. All experiments in this section were ran on a Intel(R) Xeon(R) CPU E5-2680 v3 @ 2.50GHz machine with 512GB memory. We measured running time and peak memory footprint as described in the previous section.

First we constructed *r*-index and Bowtie indexes on successively larger chromosome 19 collections (Figure 3(a), 3(b)). The *r*-index’s peak memory is substantially smaller than Bowtie’s for larger collections, and the gap grows with the collection size. At 250 chr19s, the *r*-index procedure takes about 2% of the time and 6% the peak memory of Bowtie ‘ s procedure. Bowtie fails to construct collections of more than 250 sequences due to memory exhaustion.

**Fig. 3:**
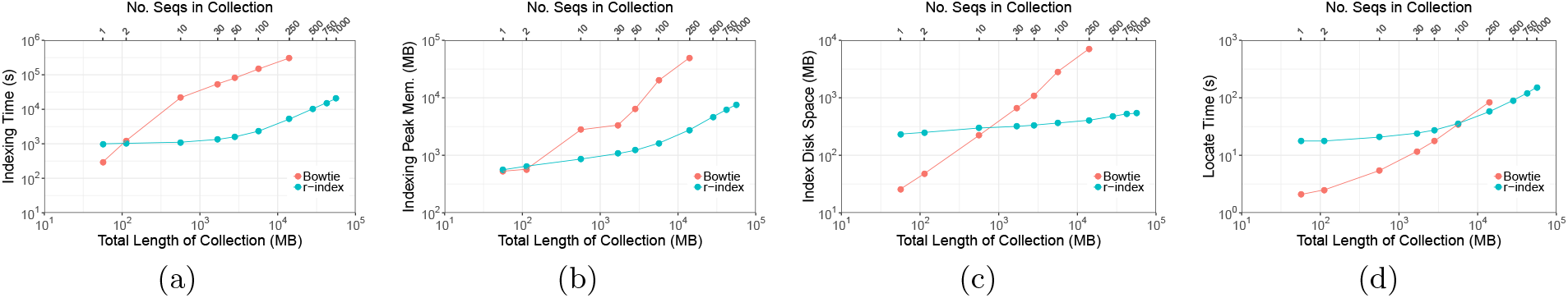
Scalability of r-index and bowtie indexes against chr19 haplotype collection size and total sequence length (megabases) with respect to index construction time (seconds) (a), index construction peak memory (megabytes) (b), index disk space (megabytes) (c), and locate time (seconds) of 100,000 100bp queries (d).

Next, we compared the disk footprint of the index files produced by Bowtie and *r*-index (Figure 3(c)). The *r*-index currently stores only the forward strand of the sequence, while the Bowtie index stores both the forward sequence and its reverse as needed by its double-indexing heuristic [19]. Since the heuristic is relevant only for approximate matching, we omit the reverse sequence in these size comparisons. We also omit the 2-bit encoding of the original text (in the `*.3.ebwt` and `*.4.ebwt` files) as these too are used only for approximate matching. Specifically, the Bowtie index size was calculated by adding the sizes of the forward `*.1.ebwt` and `*.2.ebwt` files, which contain the BWT, SA sample, and auxiliary data structures for the forward sequence. The size of the *r*-index increased more slowly than Bowtie’s, though the *r*-index was larger for the smallest collections. This is because, unlike Bowtie which samples a constant fraction of the SA elements (every 32nd by default), the density of the *r*-index SA sample depends on the ratio *n*/*r*. When the collection is small, *n*/*r* is small and more SA samples must be stored per base. At 250 sequences, the *r*-index index takes 6% the space of the Bowtie index.

We then compared the speed of the locate query for *r*-index and Bowtie. We extracted 100,000 100-character substrings from the chr19 collection of size 1, which is also contained in all larger collections. We queried these against both the Bowtie and *r*-indexes. We used the `--max-hits` option for *r*-index and the `-k` option for Bowtie to set the maximum number of hits reported to be equal to the collection size. The actual number of hits reported will often equal this number, but could be smaller (if the substring differs between individuals due to genetic variation) or larger (if the substring is from a repetitive portion of the genome). Since the source of the substrings is present in all the collections, every query is guaranteed to match at least once. As seen in Figure 3(d), the *r*-index locate query was faster for the collection of 250 chr19s. No comparison was possible for larger collections because Bowtie could not build the indexes.

### 5.2 Indexing whole human genomes

Lastly, we used *r*-index to index many human genomes at once. We repeated our measurements for successively larger collections of (concatenated) genomes. Thus, we first evaluated a series of haplotypes extracted from the 1000 Genomes Project [4] phase 3 callset (1KG). These collections ranged from 1 up to 10 genomes. As the first genome, we selected the GRCh37 reference itself. For the remaining 9, we used `bcftools consensus` to insert SNVs and other variants called by the 1000 Genomes Project for a single haplotype into the GRCh37 reference.

Second, we evaluated a series of whole-human genome assemblies from 6 different long-read assembly projects (“LRA”). We selected GRCh37 reference as the first genome, so that the first data point would coincide with that of the previous series. We then added long-read assemblies from a Chinese genome assembly project [28], a Korean genome assembly project [16] a project to assemble the well-studied NA12878 individual [15], a hydatidiform mole (known as CHM1) assembly project [30] and the Celera human genome project [20]. Compared to the series with only 1000 Genomes Project individuals, this series allowed us to measure scaling while capturing a wider range of genetic variation between humans. This is important since *de novo* human assembly projects regularly produce assemblies that differ from the human genome reference by megabases of sequence (12 megabases in the case of the Chinese assembly [28]), likely due to prevalent but hard-to-profile large-scale structural variation. Such variation was not comprehensively profiled in the 1000 Genomes Project, which relied on short reads.

The 1KG and LRA series were evaluated twice, once on the forward genome sequences and once on both the forward and reverse-complement sequences. This accounts for the fact that different *de novo* assemblies make different decisions about how to orient contigs. The *r*-index method achieves compression only with respect to the forward-oriented versions of the sequences indexed. That is, if two contigs are reverse complements of each other but otherwise identical, *r*-index achieves less compression than if their orientations matched. A more practical approach would be to index both forward and reverse-complement sequences, as Bowtie 2 [18] and BWA [22] do.

We measured the peak memory footprint when indexing these collections (Figure 4). We ran these experiments on an Intel(R) Xeon(R) CPU E5-2650 v4 @ 2.20GHz system with 256GB memory. Memory footprints for LRA grew more quickly than those for 1KG. This was expected due to the greater genetic diversity captured in the assemblies. This may also be due in part to the presence of sequencing errors in the long-read assembles; long-read technologies are more prone to indel errors than short-read technologies, for examples, and some may survive in the assemblies. Also as expected, memory footprints for the LRA series that included both forward and reverse complement sequences grew more slowly than when just the forward sequence was included. This is due to sequences that differ only (or primarily) in their orientation between assemblies. All series exhibit sublinear trends, highlighting the efficacy of *r*-index compression even when indexing genetically diverse whole-genome assemblies. Indexing the forward and reverse complement strands of 10 1KG individuals took about 6 hours and 20 minutes and the final index size was 36GB.

**Fig. 4:**
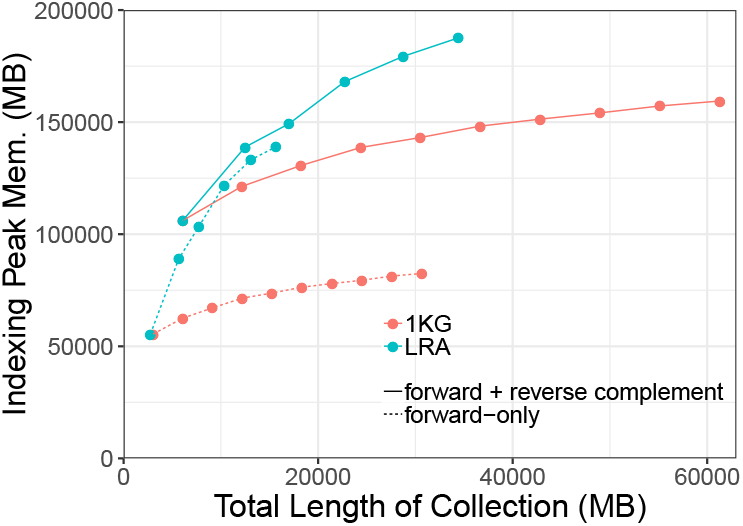
Peak index-building memory for *r*-index when indexing successively large collections of 1000-Genomes individuals (1KG) and long-read whole-genome assemblies (LRA).

We also measured lengths and *n*/*r* ratios for each collection of whole genomes (Table 1). Consistent with the memory-scaling results, we see that the *n*/*r* ratios are somewhat lower for the LRA series than for the 1KG series, likely due to greater genetic diversity in the assemblies.

**Table 1:**
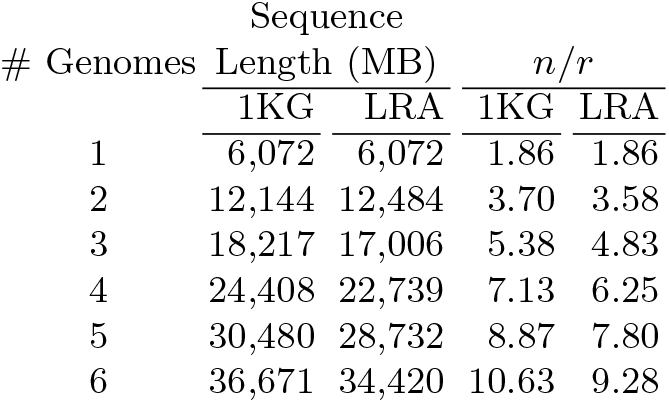
Sequence length and *n*/*r* statistic with respect to number of whole genomes for the first 6 collections in the 1000 Genomes (1KG) and long-read assembly (LRA) series.

| # Genomes | Sequence Length (MB) | | $n/r$ | |
| --- | --- | --- | --- | --- |
|  | 1KG | LRA | 1KG | LRA |
| 1 | 6,072 | 6,072 | 1.86 | 1.86 |
| 2 | 12,144 | 12,484 | 3.70 | 3.58 |
| 3 | 18,217 | 17,006 | 5.38 | 4.83 |
| 4 | 24,408 | 22,739 | 7.13 | 6.25 |
| 5 | 30,480 | 28,732 | 8.87 | 7.80 |
| 6 | 36,671 | 34,420 | 10.63 | 9.28 |

## 6 Conclusions and Future Work

We give an algorithm for building the SA and SA sample from the prefix-free parse of an input string S, which fully completes the practical challenge of building the index proposed by Gagie et al. [11]. This leads to a mechanism for building a complete index of large databases — which is the linchpin in developing practical means for pan-genomics short read alignment. In fact, we apply our method for indexing partial and whole human genomes, and show that it scales better than Bowtie with respect to both memory and time. This allows for an index to be constructed for large collections of chromosome 19s (500 or more); a task that is out of reach of Bowtie — as exceeded our limit of 512 GB of memory.

Even though this work opens up doors to indexing large collections of genomes, it also highlights problems that warrant further investigation. For example, there still remains a significant amount of work in adapting the index to work well on large sets of sequence reads. This problem not only requires the construction of the *r*-index but also an efficient means to update the index as new datasets become available. Moreover, there is interest in supporting more sophisticated queries than just pattern matching, which would allow for more complex searches of large databases.

## Footnotes

1 With the SA sample of Gagie et al. [11], this index is termed the *r*-index.

2 Given a sequence (string) *S*[1, *n*] over an alphabet *Σ* = {1,…, *σ*}, a character *c* ∈ *Σ*, and an integer *i*, rank_*c*_(*S, i*) is the number of times that c appears in *S*[1, *i*].

3 *Sampled* means that only some fraction of entries of the suffix array are stored.

4 For technical reasons, the string *S* must terminate with *w* copies of lexicographically least $ symbol.

5 Our implementation of the algorithm in Section 3, available here: https://gitlab.com/manzai/Big-BWT.

